# HiPiler: Visual Exploration of Large Genome Interaction Matrices with Interactive Small Multiples

**DOI:** 10.1101/123588

**Authors:** Fritz Lekschas, Benjamin Bach, Peter Kerpedjiev, Nils Gehlenborg, Hanspeter Pfister

**Affiliations:** Harvard University.,,.; Harvard Medical School.,.

**Keywords:** Interactive Small Multiples, Matrix Comparison, Biomedical Visualization, Genomics

## Abstract

This paper presents an interactive visualization interface—HiPiler—for the exploration and visualization of regions-of-interest in large genome interaction matrices. Genome interaction matrices approximate the physical distance of pairs of regions on the genome to each other and can contain up to 3 million rows and columns with many sparse regions. *Regions of interest* (ROIs) can be defined, e.g., by sets of adjacent rows and columns, or by specific visual patterns in the matrix. However, traditional matrix aggregation or pan-and-zoom interfaces fail in supporting search, inspection, and comparison of ROIs in such large matrices. In HiPiler, ROIs are first-class objects, represented as thumbnail-like “snippets”. Snippets can be interactively explored and grouped or laid out automatically in scatterplots, or through dimension reduction methods. Snippets are linked to the entire navigable genome interaction matrix through brushing and linking. The design of HiPiler is based on a series of semi-structured interviews with 10 domain experts involved in the analysis and interpretation of genome interaction matrices. We describe six exploration tasks that are crucial for analysis of interaction matrices and demonstrate how HiPiler supports these tasks. We report on a user study with a series of data exploration sessions with domain experts to assess the usability of HiPiler as well as to demonstrate respective findings in the data.

## 1 INTRODUCTION

The human genome is about 2 meters long and tightly folded into each cell nucleus. This results in a dense, fractal-like and three-dimensional structure in which genome sequences that are distant on the genome, can be in close spatial proximity. It has been shown [20] that this 3D structure is an important factor for regulation of gene expression, replication, DNA repair, and other biological functions. Biologists are interested in uncovering the mechanisms that drive global and local folding to better understand the vast and complex gene regulation network. This aids comprehension of the functional diversity of cells and how changes in the spatial conformation of the genome can cause diseases [24, 32, 40].

The probability of two sequences being in close proximity to each other, i.e. *interacting*, can be inferred using modern genome sequencing techniques, which yield for every genome a huge symmetric *genome interaction matrix* with up to 3 million rows and 3 million columns. Each of the 9 trillion matrix cells represents the proximity of two genomic regions. Repetitive and hierarchically nested visual patterns can be identified across the matrix, which represent so called *regions of interest* (ROIs). These patterns appear at different scales and range from hundreds of millions down to a few thousand base pairs in size.

Exploring an entire genome interaction matrix of this size to find and compare patterns of interest would require many days of work. Hence algorithms for automatic pattern extraction are being development. However, these algorithms can be very complex and often identify tens of thousands of specific pattern instances, many of questionable quality. Results of algorithms designed to identify the same type of pattern often differ substantially [18] and the lack of a ground-truth pattern collection hinders the evaluation of these algorithms. Thus, even if patterns can be retrieved automatically, assessing pattern quality requires human inspection. Moreover, interpretation of these patterns requires an informed and thorough exploration of the thousands of identified locations. In other words, data can be filtered and reduced dramatically using algorithms, but the patterns at the identified locations are still too unreliable to be visualized and analyzed further without manual inspection and evaluation.

Interactive visualization tools have been developed [45] but are focused on supporting visualization of a single or a small number of views of the matrix and navigation through pan and zoom [14, 26]. However, detailed exploration and comparison of thousands of small ROIs is unsupported by current tools yet needed, due to the size and multi-scale nature of the folded genome. In this paper, we present HiPiler—an interactive visualization tool designed for exploration and analysis of thousands of ROIs extracted from one or more genome interaction matrices (Fig. 1).

**Fig. 1.**
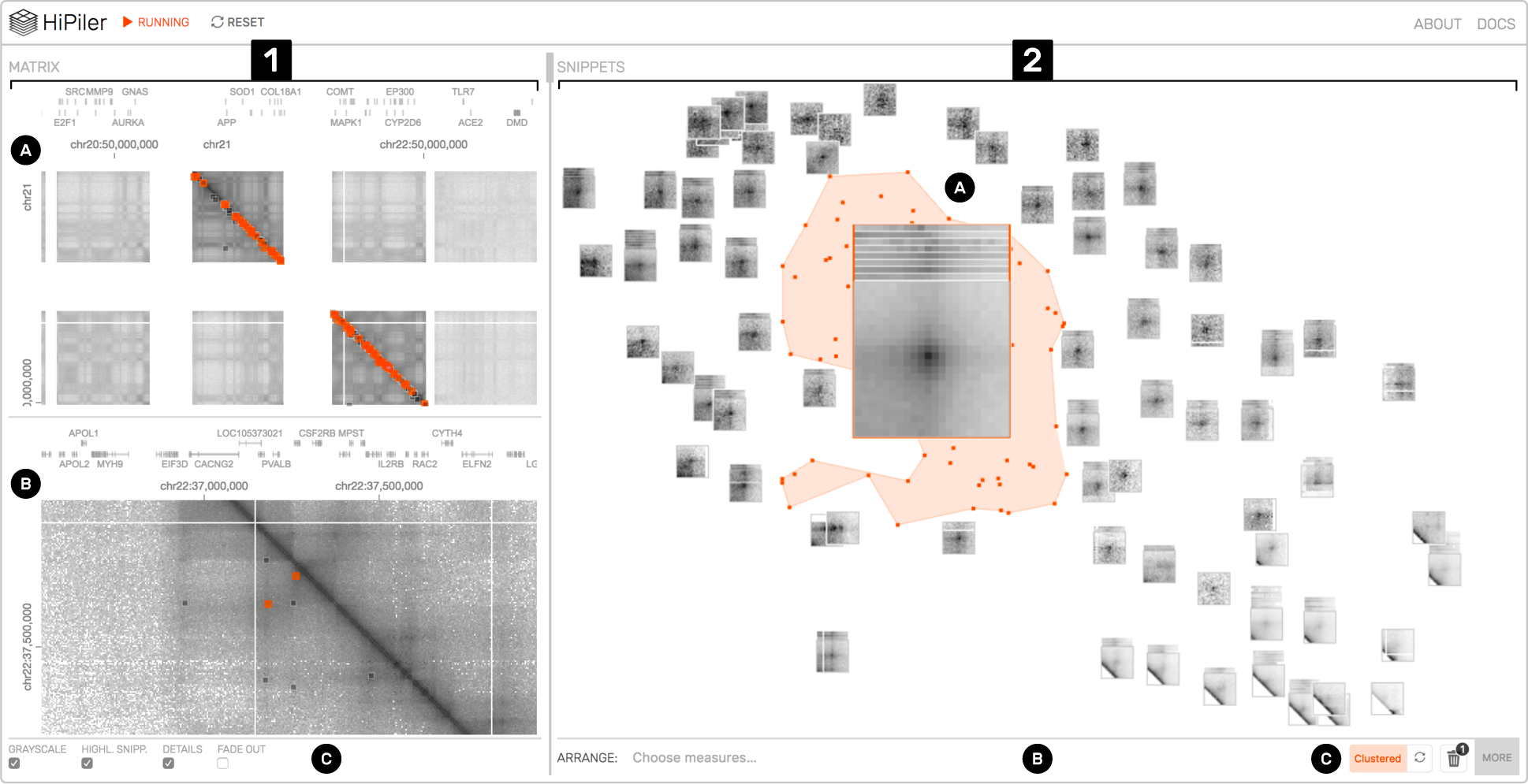
HiPiler interface: the matrix view (1) with an overview (1A) and detail (1B) matrix. The snippet view (2) presents regions of the matrix as interactive small multiples. In this example, snippets are arranged with t-SNE (2C) and a well-pronounced pile of snippets is highlighted (2A). View menus for operation are located at the bottom (1C and 2B).

To overcome the contextual constraints of exploring local patterns in very large matrices, HiPiler follows a *divide and explore* approach that extracts ROIs from the matrix and enables independent exploration (Fig. 2). HiPiler assumes a given set of ROIs, derived from specialized pattern recognition algorithms (Sect. 2.1). HiPiler then visualizes these ROIs as small heatmaps (matrices) which we call *snippets*. A snippet is associated with a set of ordinal and categorical attributes, such as its noisiness, size, or source dataset. These data are derived from the matrix itself or point to prior knowledge. Based on this data, HiPiler enables automatic and manual ordering, positioning, grouping, filtering, and visual manipulation to identify patterns present across the set of snippets (Fig. 2). Additionally, the context of snippets in the matrix is maintained through highlighting of snippet locations in the interaction matrix.

**Fig. 2.**
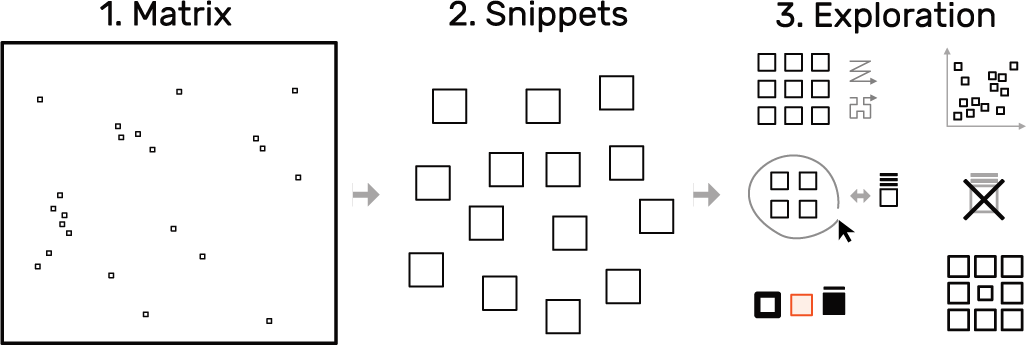
The *snippets* approach: decompose a large matrix (left) into small snippets (middle) and explore these snippets (right) using different layouts, arrangements, and styles, while maintaining the global context. The small squares within the matrix represent snippet locations.

Our design of HiPiler is informed by semi-structured interviews with ten domain experts from various genomics research labs as well as iterative design sessions over the course of several months. The interviews led to the formulation of six generic and crucial tasks for the exploration of large interaction matrices and ROIs. HiPiler is designed to support four types of scenarios: *i)* visual evaluation of the results of pattern detection algorithms; *ii)* characterization, aggregation, and outlier detection in large pattern collections; *iii)* comparison of ROIs across multiple matrices, e.g., to compare different datasets, experimental conditions, or extraction algorithms, and *iv)* correlation of matrix patterns with other genomic attributes, e.g., genes or protein-binding sites.

We evaluated the usability and appropriateness of HiPiler through a user study that involved interactive data exploration sessions with domain experts. The study results show that HiPiler is easy to learn and use, and that it offers important benefits to scientists who are analyzing and interpreting genome interaction matrices. We conclude with a list of insights and findings from the data exploration sessions, provide a list of requested features that were out of scope for this research, and outline future extensions as well as possible generalizations of our approach.

## 2 BACKGROUND—VISUAL ANALYSIS IN SPATIAL GENOME ORGANIZATION

### 2.1 Hi-C Matrix Analysis

Hi-C [31] is a method to capture genome-wide interactions. It is derived from chromosome conformation capture (3C) [11], in which genome segments cross-link when they are spatially close to one another. These cross-linked fragments are extracted, amplified, and mapped to the genome to quantify the interaction per locus (Fig. 3). Except for single-cell Hi-C experiments, a population of millions of different cells is examined at once, producing an average map of contact probabilities. Hi-C datasets are very sparse and the measured contact probabilities follow a power law decay, i.e., regions that are close to each other on the genome sequence are very likely to be in close contact while regions that are distant from each other on the genome, are expected to have almost no contact.

**Fig. 3.**
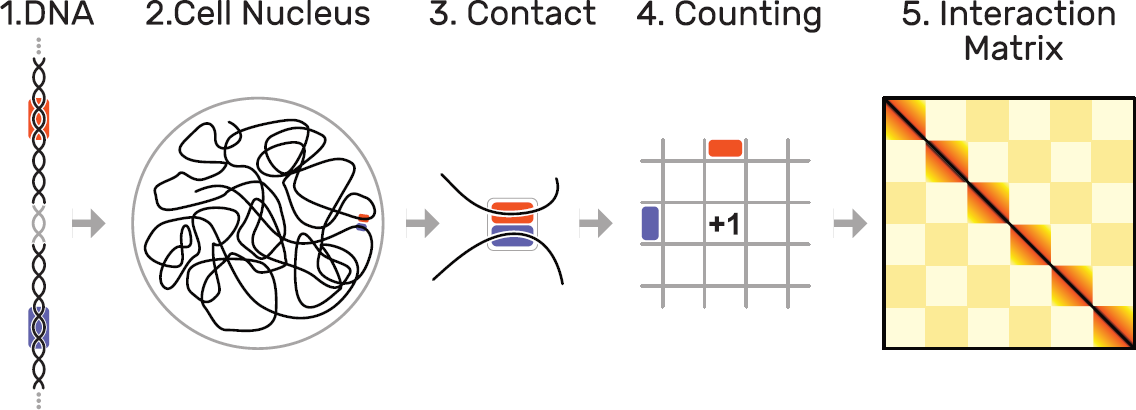
Hi-C methodology: as the DNA (1) is organized non-arbitrarily in the cell nucleus (2), certain parts (highlighted in orange and blue) are frequently in close contact (3). These contacts are quantified over a set of several hundred million cells(4), leading to interaction matrix of up to 3 by 3 million cells (5). Dark colors indicate more frequent contacts occurrences of two loci.

Despite some breakthroughs, the exact mechanisms that govern the folding of DNA are still unknown. To gain a better understanding, experts typically visualize interaction matrices as a heatmap (Fig. 1.1). Each square represents two genomic locations and the color indicates the contact probability between these regions, where darker colors represent higher probabilities that a pair of genomic regions are in close contact. Since Hi-C data offers limited insight when viewed on its own, other genomic and epigenomic data are often integrated and displayed alongside the contact probabilities. Experts usually start exploring the interaction matrix from two angles: (i) finding global patterns or visually confirming computationally-determined patterns and (ii) inspecting various patterns across regions of interest (ROIs) to identify changes under varying conditions.

Biologists define ROIs in various different ways depending on the questions being studied. Some ROIs are directly extracted by pattern recognition algorithms that work on interaction matrices [18]. In addition, other measures that are derived from different data types, such as protein-binding probability or gene expression, or existing metadata, such as genes or structural variants, are used to define ROIs too. For example, in our user study (Sect. 7) we compiled ROIs from pattern recognition algorithms for interaction matrices, from protein-binding sites, and from genomic structural variants. In all cases, extracting these ROIs is a complex problem on its own and is highly specific to the biological question. Therefore, HiPiler relies on the domain expert to compile the list of ROIs to be explored.

### 2.2 Expert Interviews

In order to identify the current challenges in the analysis of interaction matrices, we conducted a series of semi-structured interviews with ten domain experts (seven postdoctoral researchers and three graduate students). Six of the experts are computer scientists who work on algorithmic tools and pipelines. The other four experts are biologists who mainly focus on analysis of interaction matrices. Each interview lasted one hour and focused on three main parts: (i) long-term goals of genome folding-related research, (ii) workflows and strategies to gain insights, and (iii) current challenges.

The long-term goals for use of genome interaction matrices are to better understand the role of the structural organization of the genome in regards to gene regulatory and other biological processes. In this context, Hi-C analysis is also seen as a complement to existing epigenomic data. Therefore, researchers want to compare multiple conditions or subjects and use the interaction matrix to drive exploration, confirm algorithms, present findings, and generate new ideas. One of the major challenges is the size of the interaction matrix and the high number of relatively small and sparsely distributed ROIs. For example, (i) exploration of long-range interactions is cumbersome with current tools as the context is quickly lost with pan and zoom interactions. Also, (ii) the large number of pattern instances makes it hard to spot subtle differences or outliers. Finally, (iii) the data is very noisy as the folding of the genome is dynamic and not every visual pattern highlights a biological feature but could instead be caused by spatial constraints. Thus, findings needs to be verified by a number of other genomic measures as corroborating evidence.

During the interviews we learned that domain experts are comfortable with the heatmap visualization of the interaction matrices. Although the field of genome interaction matrix analysis is not mature yet, the number of well-studied patterns that correlate to biological features is limited and there are no community-defined analysis standards yet.

Several domain experts mentioned the importance of visualization for their research. For example, one expert stated that p-values are far less important to foster confidence in novel findings compared to visualization of the related matrix patterns. According to another domain expert, when developing feature extraction algorithms for interaction matrices, bioinformaticians try to model what they are seeing with their eyes. The great power of Hi-C comes from the ability to work on averages over millions of cells. At the same time, experts need to carefully check for outliers and false positives to not be fooled by the average.

### 2.3 Common Hi-C Matrix Patterns

Some of the most common patterns in the analysis of interaction matrices are shown in Fig. 4. *Loop* patterns appear as dark dots in the center and can be seen as an actual loop (Fig. 3) of the DNA. *Domains* are darker rectangles that indicate higher intra-domain than extra-domain contacts. They can be thought of as coils of DNA and are often enclosed by loops. We call loops and domain boundaries (i.e., the location where two domains meet) *point-based* patterns as they normally only span a very limited number of matrix cells. Domains as a whole can be of different sizes and are assumed to be organized hierarchically. *Flames* are horizontal or vertical streaks of darker colors that indicate higher contact probability of one locus with several others. Finally, a well-studied global phenomenon is the *checkerboard* pattern, indicating a fairly strict categorization of the genome into active and inactive compartments with high intra- and low inter-compartment contact probabilities [31]. More details on the specific biological background of the presentation patterns are described in several reviews [9, 10, 19].

**Fig. 4.**
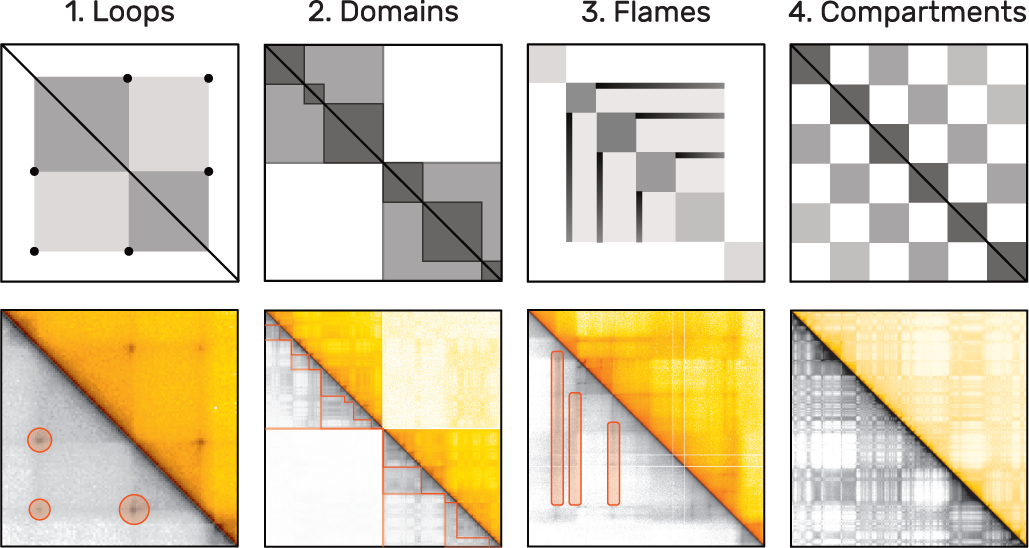
Examples of frequent patterns in interaction matrices by increasing size. The upper plots show schematic illustrations of actual examples below, taken from Rao et al. [37]. As interaction matrices are symmetric, the lower triangular matrix is displayed in grayscale to highlight the patterns using orange markers. *Loops* (1) appear as dark central dots and span only few cells. *Domains* (2) are darker rectangles that are presumably organized hierarchically. *Flames* (3) are horizontal or vertical lines. Active and inactive compartments of the genome create a global *checkerboard* pattern (4).

### 2.4 Hi-C Matrix-Analysis Tasks

Based on our interviews, we identified six tasks related to the exploration of interaction genome matrices. We generalized these tasks to pattern exploration in other large matrices (boldface text) and provide the rationale in Hi-C analysis (normal text):

**T1 Search for known patterns:** Some visual patterns are known to have a specific biological meaning. Experts typically look for them first.
**T2 Discover new patterns:** When studying multiple ROIs, experts often find a variety of recurrent patterns, which have no known biological functions.
**T3 Study one instance of a pattern:** Once a pattern or ROI has been identified, domain experts are interested in studying the details of this pattern.
**T4 Compare instances of one pattern type:** The variance of a pattern types and their distributions in interaction matrices are essential for studying biological features.
**T5 Correlate pattern instances with features:** Snippets are associated with additional attributes describing other genomic features. Experts want to identify correlations between these attributes and the patterns.
**T6 Compare ROIs across matrices:** Experts want to compare ROI across multiple matrices (e.g., different experimental settings or replicates) to draw causal relationships and assess the stability of patterns.

## 3 RELATED WORK

### Visualizing Genome Interaction Matrices

Advances in high-throughput DNA sequencing have led to a notable increase of available interaction matrices. This sparked development of several specialized software tools for visualization [45]. All of these tools visualize the data in the form of a large matrix, with contact probabilities being translated into color maps, and support pan-and-zoom as a means of navigation. Most applications work offline and integrate 1D tracks, which show various genomic measures in the forms of line graphs or bar charts. HiGlass [26] is a web-based interaction matrix and 1D track viewer that additionally provides seamless view manipulation and view sharing. We integrated HiGlass into HiPiler to provide an overview of the snippet locations, to display 1D tracks, and to select and highlight snippets from the interaction matrix. Pattern-centric visualizations of interaction matrices are currently not supported by any tool. Also, experts use Matplotlib [25] to visualize aggregations of patterns in multiple different ROIs or experimental conditions.

### Visualizing Large Matrices

Matrices are a common representation for visualizing networks or graphs [6, 43]. Thus, we briefly overview related visualization techniques that focus on large matrices. Large matrices make it hard to analyze detailed visual patterns and to compare distant parts in the matrix. Common interactions, such as panning and zooming, are *content-agnostic* as they operate entirely on the view level but do not perform any operations on the data itself.

On the other hand, *content-aware* approaches incorporate the data to drive visualization. For example, ZAME [16] aggregates individual cells into higher-level cells as the user zooms out. Zoomed-out cells then show a glyph with the distribution of values grouped by this cell. Such zoom-based interfaces can scale well for very large matrices, but they loose fine-grained visual patterns that are required in interaction matrix analysis. Rather than aggregating cells, Melange [17] allows skipping rows and cells by literally folding them into a third dimension with the effect of reducing the size of the visible matrix. Hence, the remaining non-folded parts are observed in more detail and visual patterns can be compared across long distances. Dinkla et al. [12] present a technique for visually compressing gene regulatory networks and which takes the underlying data into account. The compressed representation works for medium-sized networks but still becomes overwhelmingly large as the network grows. NodeTrix [23] is a hybrid approach that shows only clusters as matrices and visualizes connections between clusters as straight lines.

While all these approaches improve exploration of large matrices, our approach is different in that we do not study exploration of the entire matrix but focus on a set of ROIs of the matrix (snippets). This approach provides enough expressive power to focus on important parts in the matrix only and makes visual analysis dependent on the number of ROIs instead of the actual size of the matrix.

### Divide-and-Conquer Approaches

Mining specific patterns in large datasets for display and exploration has been employed successfully in other domains, e.g., image and network analysis. Network motifs [35] refer to subgraphs in a network and are similar to our matrix snippets. Network motifs have frequently been used as first-class objects in network exploration. Visualized in an adjacency matrix, they result in recurring visual patterns [6]. For example, clusters occur as rectangles (similar to *domains in interaction matrices*) and highly connected nodes appear as horizontal and vertical lines (similar to *flames*). Schreiber et al. [39] allow for the selection of a sub-graph (network motif) in a network and consequently retrieve and visualize all occurrences of sub-graphs with similar network topology. Dunne and Shneiderman [13] first detect network motifs such as clusters and fans and then replace their occurrence by specialized glyphs in the node-link diagram of the network. Von Landesberger et al. [42] extract motifs, obtain metrics (e.g., size, density, average degree and others), and visualize the retrieved motifs using a self-organizing-map [28] layout. Except for Cubix [4], which visualizes the evolution of ego-networks through heatmaps similar to adjacency matrices, network motif visualization has so far been focused on node-link representations, and not been seen as an approach for interactive visualization of large matrices. While the approach of von Landesberger et al. [42] could be generalized to adjacency matrices, explicit interactive means for exploration and alternative motif layouts were not the focus of their work. While we can derive inspiration from these approaches, snippets from interaction matrices are very different from network motifs with respect to visual appearance.

Our work is most inspired by MultiPiles [3]—an interface that employs the visual and interactive metaphor of piling adjacency matrices and exploring these piles to visualize time sequences in dynamic networks. We integrate the piling metaphor and some of the exploration features from MultiPiles but heavily extend upon them by introducing linear ordering, multi-dimensional arrangements, clustering, filtering, and grouping approaches for exploring many snippets.

## 4 DESIGN OF HIPILER

The key goal for HiPiler is to enable the exploration of many ROIs in a large matrix via interactive small multiples that represent snippets. Snippets can be ordered, arranged, filtered, and grouped independently of their neighborhood (Fig. 2). Extracting ROIs from the interaction matrix is not part of HiPiler, instead we assume they are given, either resulting from manual search or an algorithm. Snippets can represent differently sized ROIs with patterns of different shape but need to have the same aspect ratio to be scalable to an equal screen size.

Design and development of HiPiler was driven by questions such as *“How can we meaningfully limit the number of snippets shown?”*, *“Which interactions are important to efficiently support arrangements?”*, or *“How to effectively link the interaction matrix with the snippets”*. In the following sections we will describe the data model, design, and interactions of HiPiler that are guided by the identified tasks (Sect. 2.4).

HiPiler consists of two main views: the matrix view and the snippets view (Fig. 1.1 and Fig. 1.2 respectively). The matrix view displays one or more interaction matrices and supports two display modes: *overview* and *detail*. The upper half of the matrix view contains the overview (Fig. 1.1A) and indicates the location of all explored snippets. The detail matrix (Fig. 1.1B) is in the lower half and enables browsing the interaction matrix via pan-and-zoom. Both matrices highlight the snippet location with colored rectangles. A menu at the bottom (Fig. 1.1C) provides options for customization. The snippet view (Fig. 1.2) displays snippets according to user-defined ordering, arrangements, filtering, or grouping. For example, in Fig. 1.2 snippets have been arranged with t-SNE [33] (Fig. 1.2C). The center of the plot features a scaled-up pile of multiple snippets (Fig. 1.2A). The original location of the piled-up snippets is indicated by the orange hull drawn in the background. Other means of operating the snippet view are provided via a menu at the bottom (Fig. 1.2C). An overview of all conceptual design aspects of HiPiler is given in Fig. 5.

**Fig. 5.**
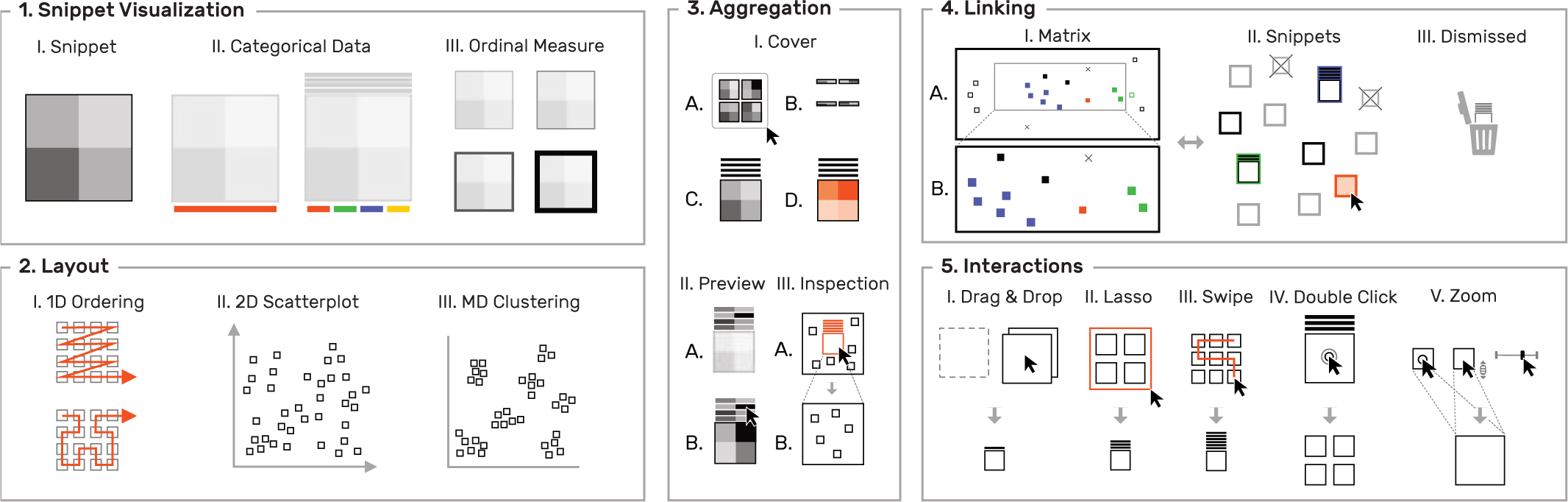
Overview of visualization and interaction concepts in HiPiler.

### 4.1 Data Model

Many graph and network data can be represented as a matrix, where the <i,j> cell contains a correlation measure between node i and node j. In addition, cells can be associated with multiple categorical and ordinal attributes. These attributes can be measures or annotations derived from the matrix or given by prior knowledge. For example, noise, pattern sharpness, or distance-to-diagonal are derived from the matrix and referred to as *data* attributes. Prior knowledge, which we refer to as *meta* attributes, can for example be confidence in the correlation measure, protein-binding sites, or gene expression levels. HiPiler assumes a fixed ordering of rows and columns, which in our case is given by the genome sequence. Each ROI is defined as a set of start and end locations and we assume, but do not require, that datasets contain many small ROIs that are distributed across the interaction matrix.

### 4.2 Design Aspects

During the development of a prototypical implementation, we met with three of the initial and one additional domain expert (all are computational biologists) for 1-2 hours to collect feedback on our design choices. We additionally presented the prototype to a group of domain experts that haven’t been involved in the initial interviews to get unbiased feedback. We explored different visual designs, such as node-link or arc diagrams, but given the dominance of the matrix visualization for genomic interaction matrices, the mixed feedback of experts regarding alternative representations (Sect. 2.2), and the generality of matrix visualizations, we focused on matrix techniques. Most important, we think matrix snippets are best to show the respective patterns visually.

**Snippet Metaphor:** Snippets are the essential building blocks of HiPiler as they help experts to identify known (T1) and unknown patterns (T2) among ROIs. In addition to displaying a part of the matrix, snippets can be associated with categorical and ordinal attributes, which are displayed with additional visual marks (Fig. 6); addressing T3 and T5. The visualization of these attributes has been kept minimal to act as information scent [36] and to avoid distraction from the snippets.

**Fig. 6.**
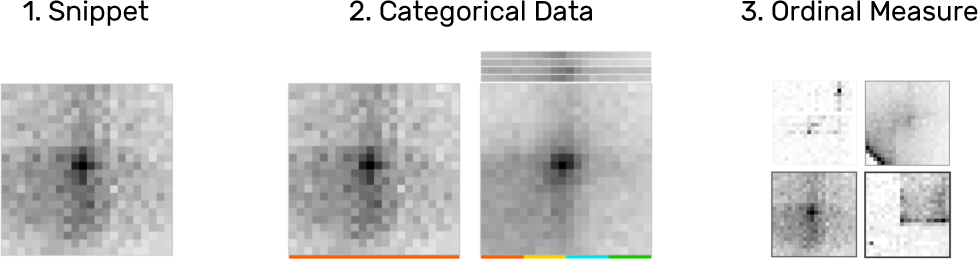
Snippets are drawn as a heatmap (1), showing the matrix data of their ROI. Categorical attributes can be visualized with color tags (2). An ordinal measure associated with a snippet can be shown via frame width and color (3). For example, a high value is drawn as a dark thick border.

**Snippet Layout:** Domain experts want to explore several hundreds to thousands of instances of one pattern type simultaneously (**T4**) to identify groups and uncover potential correlations between patterns in the interaction matrix and other genomic features. To support these tasks, HiPiler’s layout is entirely data and attribute-driven; allowing for one-dimensional (1D) ordering, two-dimensional (2D) arrangements, or multi-dimensional (MD) clustering via dimensionality reduction (Fig. 7).

**Fig. 7.**
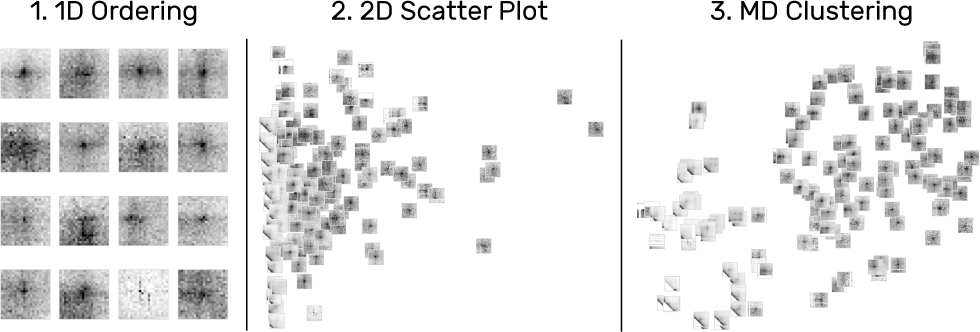
Snippets can be arrange along various different dimensions. For a single attribute, snippets are laid out in 1D supporting reading direction and Hilbert curves (1). Selecting two attributes creates a scatter plot (2). For more than two attributes HiPiler applies dimensionality reduction algorithms (3) such as t-SNE [33].

**Aggregation:** To support the exploration of large numbers of snippets, HiPiler applies and extends upon the piling metaphor of Multipiles [3]. Snippets are stacked into a pile featuring a *cover* matrix that shows a summary of the stacked snippets. The cover is calculated by taking the mean or standard deviation of all snippets (Fig. 8.1C and 8.1d). These *cover modes* help experts to assess the average expression and variance of patterns (T4 and T6). To briefly browse piled snippets, HiPiler displays up to eight *pile previews* as 1D heatmaps above the cover (Fig. 8.1A). Previews show the mean column values of their underlying matrix data. Moving the mouse cursor over a preview temporarily displays the related snippet (Fig. 8.2). For a large number of snippets per pile it would be inefficient to limit the exploration to the cover or individual (dispersed) snippets only. Therefore, we have added support for hierarchical *inspection* of piles. When inspecting a pile, only the snippets of the pile are shown and the layout is automatically scaled to accommodate the region occupied by these snippets only (Fig. 8.3).

**Fig. 8.**
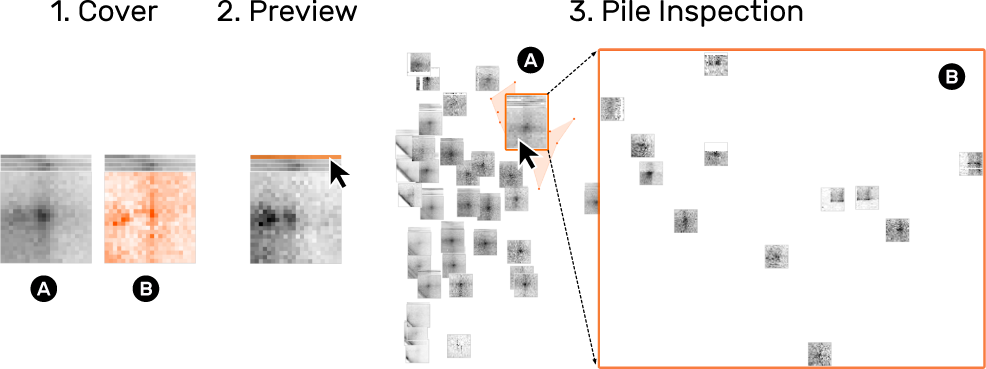
HiPiler displays a cover matrix of the average (1A) or variance (1B) of snippets on a pile. Additionally, each snippet is displayed as a 1D preview showing a horizontal aggregate of the snippet’s rows (2). Moving the mouse cursor over a preview shows the related matrix. Inspecting a pile (3B) temporarily hides all other snippets (3B).

**Fig. 9.**
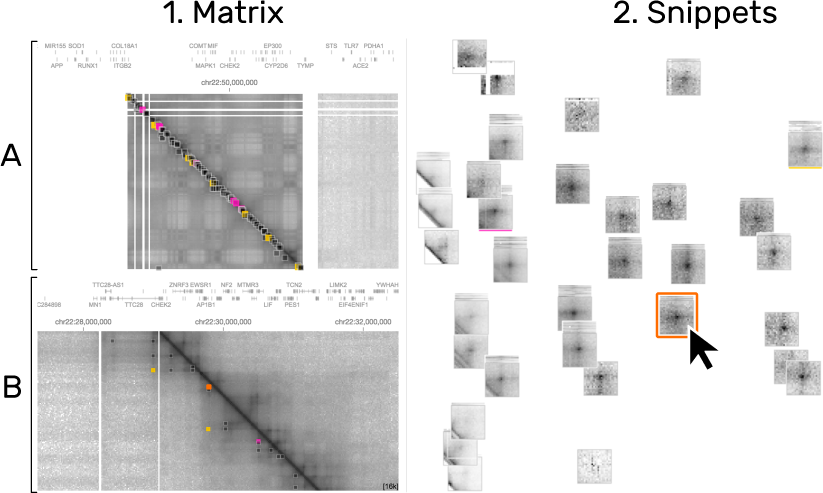
The location of the hoverd snippet are highlighted in the interaction matrix (1A) and the matrix detail (1B) in orange. Other colors (pink, orange, and yellow) represent otherwise selected matrices. HiPiler distinguishes between groups of snippets via color tags (2. pink and yellow bar) and manual highlighting (2. orange).

**Fig. 10.**
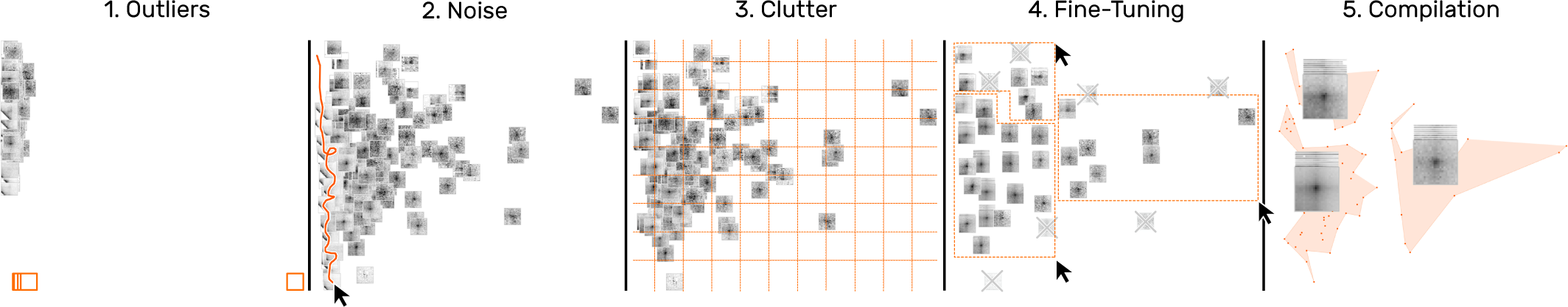
Snippet curation via filtering. A typical filtering process involves the removal of outliers (1) and noise (2), clutter reduction via automatic piling (3), and manual grouping (4) until a satisfactory collection of snippets is obtained (5).

**Linking:** In almost all cases the neighborhood of snippets is crucial as the genomic locations associated with snippets act as the *ground-truth* in genome biology. In HiPiler, snippets are therefore interconnected with the matrix view by highlighting their location via colored rectangles (Fig. 4.2.1). Snippet locations can be shown permanently via color tags or temporarily by selection. To support the investigation of the neighborhood of a snippet, HiPiler implements a *detail matrix* view (Fig. 4.2.1B). It is possible to fade out snippets that are not visible in the viewport of the detail matrix to provide more focus. Also, HiGlass [26] supports 1D genomic tracks, which enables experts to correlate patterns to many other genomic measures (T5). Finally, the matrix view can host more than one matrix to support comparison across datasets (T6).

### 4.3 User Interaction Techniques

We have adopted piling interactions from Multipiles [3], where the user can manually create piles with drag-and-drop (Fig. 5.5.I) or lasso selection (Fig. 5.5.II) and disperse piles via double-clicking (Fig. 5.5.IV). In addition, HiPiler implements more fine-grained controls for piling and zooming. For 2D and MD layouts, HiPiler automatically downscales the size of snippets and piles to fit a larger number of snippets on the screen and to avoid clutter. To make piling in dense layouts easier, we added swipe-based pile selection, where the user can move the mouse cursor over the snippets to be selected while holding down the left mouse button (Fig. 5.5.III). Also, temporary upscaling of individual snippets is supported by steering the scroll wheel while having the mouse cursor placed over the target snippet (Fig. 5.5.V). It is also possible to zoom into the entire snippet view to inspect a sub-region.

## 5 USAGE SCENARIOS

In the following usage scenarios we demonstrate how HiPiler can be used to study the diversity and variance of patterns using a set of loops previously reported by Rao et al. [37]. A perfect loop pattern exhibits a dark central dot surrounded by relatively bright areas (Fig. 1.2a). Since genome interaction data is sparse and noisy and since there are no gold standards for pattern extraction yet (Sect. 2), questions that guide our exploration are: *“How do average patterns at extracted locations look?”*, *“Can we compile a set of snippets with well-pronounced loop patterns?”*, *“Are we missing locations which express the same or similar patterns?”*, *“Is there a correlation between patterns and other attributes?”*, and *“Can we see similar patterns at the same locations in other matrices?”*.

### Overview

A common first step in studying pattern variability is to gain a quick overview of the entire set of snippets and the general diversity of patterns. After loading the data, some snippets in the snippet view with loop patterns are immediately visible (T1) (Fig. 11.1A), while others contain parts of the diagonal, are noisy, or appear to be empty (T2) (Fig. 11.1B). Ordering snippets by their distance to the diagonal uncovers more consecutive snippets with similar patterns (Fig. 11.2). Since the number of snippets is too large to get an overview, we group snippets by their pairwise Euclidean distance of the underlying data so that scrolling is avoided (Fig. 11.3). This essentially piles up snippets hierarchically into *k* clusters, where *k* is the maximum number of snippets that can be displayed at the current size so that no two snippets or piles overlap. The covers show the mean patterns of piles, indicating that some exhibit a well-pronounced loop pattern (Fig. 11.3A) while others are more diverse (Fig. 11.3B).

The default cover displays the mean signal across all snippets on a pile (Fig. 12.1). Changing the cover mode to variance shows the standard deviation of piled-up snippets and supports assessing pattern variance (T4) (Fig. 12.2). Piles with a well-pronounced mean loop pattern do not usually express significant variance, indicated by a relatively *flat* heatmap (Fig. 12.2A). The variance cover shows significantly darker and more saturated spots for piles containing noisy snippets or outliers (Fig. 12.2B).

To get a better sense of the pile composition (T4), moving the mouse cursor over snippet previews temporarily shows the previewed snippet as a whole on the cover (Fig. 12.3). HiPiler limits the overall number of previews to *i*, which is configurable, to prevent occlusion by high stacks of previews. When a pile consists of more than *i* snippets HiPiler utilizes k-means clustering [34] to group the snippets.

### Filtering

One of the initial questions involves dissecting noisy and well-pronounced patterns. Arranging snippets by noise and their distance to the diagonal transforms the 1D ordering into a 2D scatter plot. This spreads out snippets spatially (Fig. 10.1) and supports better differentiation between groups (T1 and T2). First, we notice some outliers, which are completely white and far away from the diagonal (Fig. 10.1 and 1B). Dismissing outliers (i.e., moving them to the trash) re-scales the scatter plot (Fig. 10.2). The *swipe* selection is useful for non-linear fine-grained piling of snippets in dense areas (Fig. 10.2). Moving the mouse over snippets while holding down the left mouse button leaves a trail of which snippets are to be grouped (Fig. 10.2 orange line). Next, to quickly reduce clutter one can auto-pile snippets by grid cells (Fig. 10.3). Manual piling via drag-and-drop or *lasso* selection and dismissing further supports to filter the set of snippets (Fig. 10.4) until a satisfactory collection of well-pronounced loops is obtained. Placing the mouse cursor over a pile displays the location of its piled up snippets given the current arrangement (Fig. 10.5).

**Fig. 13.**
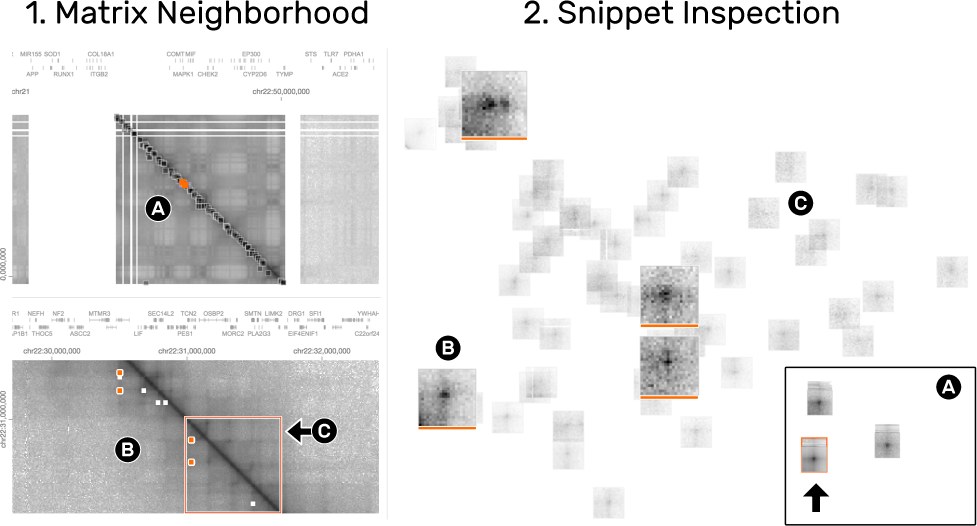
The matrix (1) and snippets (2) views are highly interconnected to enable the exploration of the spatial neighborhood of snippets. The detail matrix (1B) shows the colored snippets (2B) in context. To focus on the currently visible neighborhood, non-visible snippets are faded-out (2C).

### Snippet Neighborhood

Having curated a collection of snippets with well-pronounced loop patterns, one task is to study the distribution of the piled-up snippets across the interaction matrix. Clicking on a pile highlights the location of its snippets as orange rectangles in the matrix (Fig.1.1). *Inspecting* a pile displays only its snippets while hiding any other piles or snippets (Fig. 13.2A). We notice a region which features many highlighted snippets (Fig. 13.1A). Navigating to this region via zoom-and-pan provides spatial context to the snippets (Fig. 13.1B). Being able to explore the neighborhood of snippets is important for correlation of different pattern types, e.g., we can see that the highlighted snippets appear within a dark rectangular area, known as TADs (Sect. 2.4) (Fig. 13.1C). We also find other loop-like patterns that have not been detected by the algorithm. To identify which snippet is currently visible in the detail matrix other non-visible snippets are faded out in the snippet view (Fig. 13.2C). Color tags can be used to permanently highlight snippets (Fig. 13.2B).

**Fig. 11.**
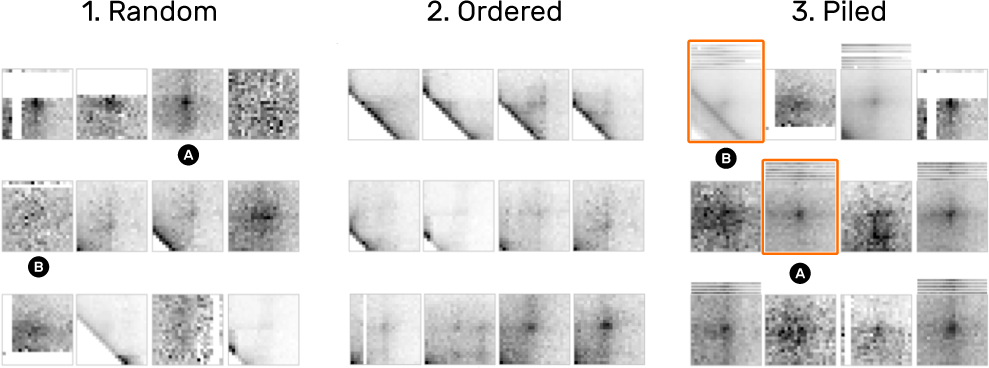
1D ordering and similarity piling. (1) Random arrangement of snippets. (2) Snippets ordered by their distance to the diagonal show a progressively emerging pattern. (3) Piling by pairwise similarity highlights similar patterns and outliers. For example, 3A shows a well-pronounced loop pattern while 3b is more noisy due to the diagonals.

**Fig. 12.**
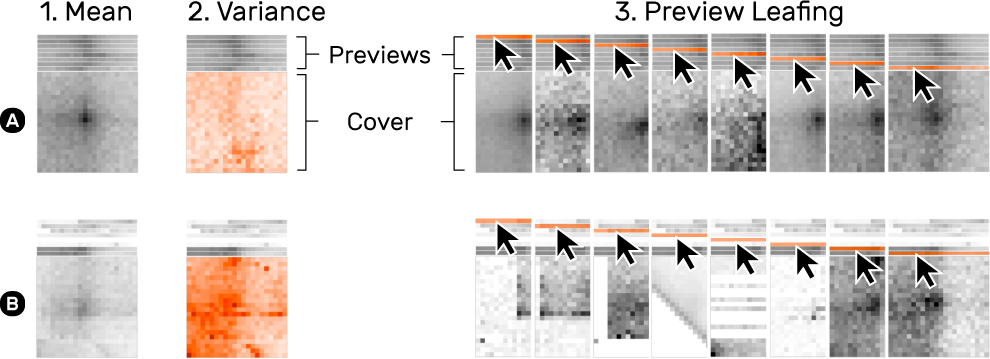
Snippet aggregation of a more homogeneous (A) and diverse (B) set of snippets. The default cover shows the mean of snippets on a pile (1). The variance cover mode (2) highlights the deviation of snippets. Moving the mouse cursor over a pile’s previews temporarily shows the corresponding snippet or pile (3).

### Correlation with other Genomic Features

HiPiler supports three ways to investigate correlations between snippets and other attributes (T5): i) integrating additional genomic tracks in the matrix view (Fig. 14.1), ii) visualizing an attribute via frame encoding (Fig. 14.2), or iii) arranging snippets according to meta attributes. For example, the previously colored snippets that show a pronounced loop pattern perfectly align with three other genomic measures displayed above the interaction matrix (Fig. 14.1A and 1B), suggesting that this ROI exhibits a biological function. Frame encoding is useful for integrating attributes into the snippets view and act as information scent [36] for further investigation.

**Fig. 14.**
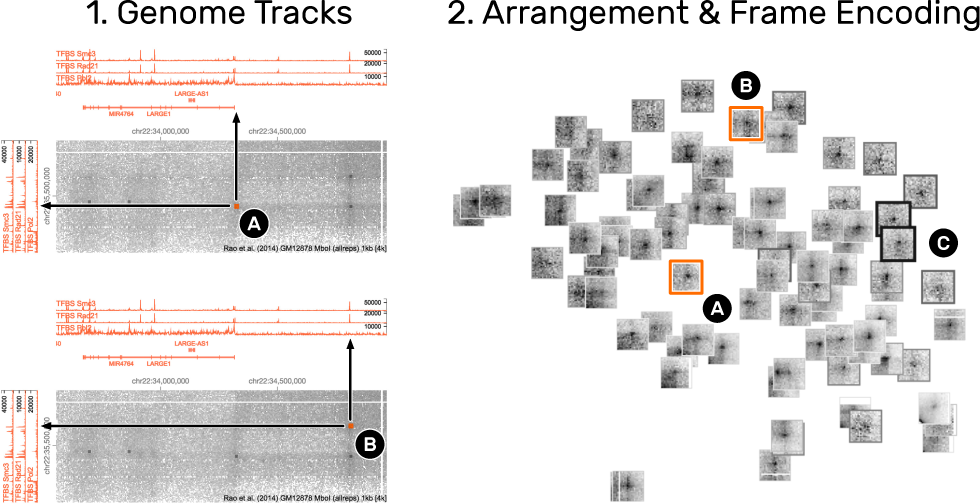
Correlation of snippets with other attributes. (1) The integration of line graphs next to the interaction matrix show genomic measures. 1A and 1B are two snippets (2A and 2B) that exhibit a loop pattern but are correlated with the expression (peaks in the line graph) of different genomic measures, indicating that the underlying ROIs serve different biological functions. Numerical measures can also be integrated into the snippets view (2) by alternating the frame width and color of snippets (2C). Thicker and darker frames indicate higher values.

### Comparison across Matrices

Comparing ROIs across matrices helps to differentiate the variance of patterns. Extracting the same ROIs in both matrices allows for pairwise comparison of snippets. *Grouping by location* is an operation that automatically piles the pairs of snippets. Activating the variance cover mode highlights the deviation of paired snippets. For example, a pronounced loop pattern uncovers high deviation around the center of the snippets (Fig. 15.2B), meaning that only one of the two snippets contains the loop pattern (Fig. 15.2C and 15.2D). Using this technique in combination with the detail matrix view allows to identify ROIs where the respective loop pattern disappears (Fig. 15.1A and 15.1B).

**Fig. 15.**
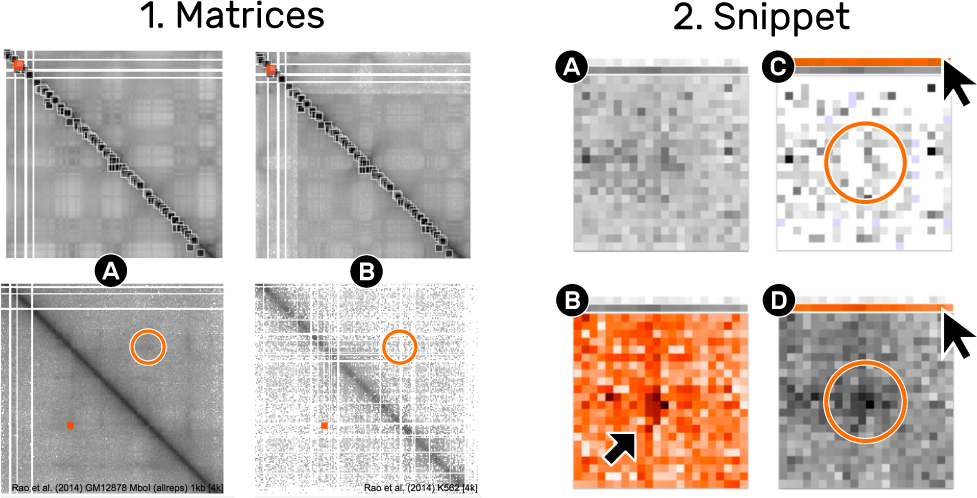
Comparison of one location in two different matrices. The matrix view (1) shows two different matrices (1A and 1B). The ROI indicated by the small square rectangle is extracted from both matrices, displayed as a pile (2A). The mean cover (2A) does not show a pronounced loop pattern in contrast to the variance cover (2B). The detail matrix view indicates that the right matrix (1b) is less dense than the left matrix. Leafing over the previews reveals that the loop pattern is completely gone in (1C) compared to (1D).

### Further Use Cases

HiPiler supports exploration of snippets of any equally-sized ROIs, for example, promoter-enhancer pairs, domain boundaries, telomeric regions, or structural variations. While the pattern types differ, the tasks and resulting interactions are essentially identical.

## 6 IMPLEMENTATION

HiPiler is implemented as a web application consisting of a front-end interface for the visualizations and a back-end server that provides the data. The front-end is entirely written in JavaScript utilizing Aurelia [15] as its application framework and Redux [2] for fine-grained, history-aware state management. The matrix snippets are visualized with WebGL using Three.js [8] as a middleware. Finally, HiGlass [27] is integrated as a library for displaying the interaction matrix and genomic tracks. The back-end serves data to HiGlass and provides the matrix snippets. The server is implemented in Python and uses Django as its application framework. The contact matrices are accessed through Cooler [1], a Python-based service library for storing and querying of Hi-C data. A custom API endpoint extracts subsets from an interaction matrix defined by two genomic ROIs and the zoom level. The front and back-end are two separate applications that can be decoupled to load different data types. HiPiler is open source and available on GitHub (https://github.com/flekschas/hipiler).

## 7 USER EVALUATION

We conducted a qualitative user study with domain experts to investigate the utility and usability of our approach. We obtained agreement from three of the domain experts that we had interviewed earlier as well as from two additional domain experts for an open-ended data exploration session with HiPiler. All experts are computational biologists that work with genome conformation contact data on a daily basis. Two of the domain experts are senior PhD students while the other three are postdoctoral researchers. Their work experience in the field ranges from one to eight years.

We conduced individual sessions that lasted between 1-2 hours. First, we introduced the core concepts of HiPiler and gave a walkthrough of its main functionalities (10-20 minutes) (Sect.4). Afterwards, each domain expert completed a training session in which they performed specific interactions, e.g., piling up snippets with lasso selection or arranging snippets by some attributes, on a training dataset for 10 minutes. Next, we loaded the expert’s own datasets and let them explore the data by themselves (30-90 min). During the exploration, we asked the experts to think aloud and to express their rationale behind operations. The entire study was performed on a pre-configured computer equipped with a 27-inch external monitor and a standard mouse. Keyboard input was not required. We recorded screen content and audio during each session for later analysis.

### 7.1 Exploratory Sessions

Participants (P) 1, 3, and 4 investigated a set of pairwise enhancer-promoter locations [41], which are assumed to interact structurally [22, 29, 30]. In the interaction matrix, structural interactions appear as dark spots, forming the patterns described in Sect. 2.3. P1, P3, and P4 started browsing the snippets to get an overview: P1 and P3 manually piled snippets that showed a loop-like pattern (T1). P1 ordered snippets by their distance to the diagonal first and then switched to dimensionality reduction with t-SNE to better differentiate between noisy and well-pronounced patterns (T4). P1 and P3 reduced the visual complexity by piling up noisy snippets using the swipe selection tool to subsequently dismiss them. P3 found a snippet with a non-centric loop pattern (Supplementary Fig. S1.1) and investigated its spatial neighborhood using the detail matrix view (T3). P3 noticed that several snippets are located in relative proximity to one another, which is shown by the colored squares in the interaction matrix. P3 decided to keep only one snippet with a strongly pronounced pattern to avoid overrepresentation (T3 and T4). Similar to P1, P3 continued with arranging snippets using t-SNE and found more loop-like patterns after examining noisy groups of snippets, indicating that some locations exhibit structural interactions (T4). P4 first checked the overall quality of snippets and noticed high sparsity within the snippets’s matrix, which results in salt-and-pepper-like noise. By activating the visualization of low quality cells, P4 found that the large number of low quality cells indicates that these noisy snippets come from a region of low quality (Supplementary Fig. S1.2). Finally, P3 opened the detail matrix view and navigated to the location of a set of snippets showing a loop-like pattern (T2). They concluded that the patterns are potentially related to another biological feature (T5) after finding additional patterns in the matrix view.

P2 explored loop patterns as reported in the literature [37] and wanted to determine the performance of a detection algorithm (T1). P2 started arranging snippets by their distance to the diagonal and noise and identify outliers (T3). To study snippets with a well-pronounced loop pattern, P2 decided to remove noisy snippets first. Finally, P2 tested the t-SNE-based snippet arrangement to further dissect noise from clean patterns and refined the dissection by iteratively applying t-SNE followed by the removal of noisy snippets (T4).

P5 studied structural variations in the genome (e.g., deletions, insertions, or translocations of DNA sequences) and wanted to assess the performance of predicted results from data analysis tools like Delly [38] or Meerkat [44] (T5). They loaded structural deletion sites that are expected to show half empty snippets (T1). Empty snippets can be the result of a i) structural variation or ii) technical limitations of the current technology for generating interaction matrices. To distinguish between them, P5 activated the visualization of low quality cells. P5 piled up and removed some empty (white) snippets and shifted their focus to unexpected non-empty snippets for detailed investigation (T2). P5 browsed the spatial neighborhood of the respective snippet in the detail matrix view and loaded a second interaction matrix for comparison (T6). The second matrix is assumed to have no or less structural variations. P5 found significantly brighter columns and rows in one of the two interaction matrices indicating a true DNA deletion (Supplementary Fig. S1.3) (T5).

A chronological summary of the participant-specific actions is provided in Supplementary Table S1.

### 7.2 Findings

**Snippets are useful** for exploring hundreds to thousands of pattern occurrences in interaction matrices. All participants stated that this technique enables them to easily assess the variety and variance of patterns. P1, P3, and P4 pointed out that seeing what an average pattern is composed of is particularly helpful to avoid misinterpretation based on the inclusion of noise or unrelated patterns. The snippets approach further aided P1, P3, and P4 to determine reasonable thresholds of attributes for the exclusion of noisy patterns, i.e., they visually determined at which value they would consider a snippet to be labeled noisy. They note that the snippet view can be used to select promising candidates for further investigation or to build a set of “ground truth” ROIs for evaluating the performance of pattern detection algorithms.

**Coordination between the matrix and snippet view is highly appreciated** by every participant as many tasks require spatial context. The interplay between the two views enables the participants to explore snippets in new ways, e.g., P2 states that HiPiler enables them to correlate patterns according to prior knowledge while still maintaining context. P1 pointed out that they usually don’t know what they can expect to see in a interaction matrix and that it is great to be able to browse the neighborhood of snippets when they spot surprising patterns. P5 noted that for their research questions it is essential to have the matrix view since some biological phenomena lead to patterns that span the entire matrix. During all sessions, the participants spent an equal amount of time on the snippets view, arranging and organizing snippets, and the matrix view, reconfirming findings and further exploring patterns in the neighborhood of snippets.

**Identified tasks shown to be valid** as all of them have been addressed at least once during the study (Supplementary Table S1). Though, the focus on specific tasks naturally depended on the respective dataset.

**Users quickly grasp the main operations** supported by HiPiler. After the guided 10 minute training session, all participants in our study were able to explore snippets on their own. All participants noted that HiPiler is very easy to learn. P2 said *“the menus are where you would expect them to be”* and P4 stated that HiPiler *“is much more thoughtful than expected”* in comparison to current visualization tools for interaction matrices. However, we acknowledge that all participants are proficient in operating computers and that an initial phase of training is necessary.

**HiPiler significantly improves on the state-of-the-art** tools for genome interaction matrix exploration. Current tools are currently limited to pan-and-zoom interactions of the entire interaction matrix or require custom, code heavy solutions with Matplotlib [25]. P4 noted *“[HiPiler] takes it from zero to infinity”* to point out that there are no other feature-centric visualization tools for interaction matrices available.

**Participants strongly indicated that they will use HiPiler** for their research once additional features are implemented. These features include data processing for HiPiler and displaying of various numerical and statistical attributes related to snippets, e.g., translating visual encodings back to their numerical values.

## 8 DISCUSSION

### Additional Features

During the user study (Sect.7) domain experts suggested a number of additional features that would make further exploration of ROIs more efficient for analyses on a daily basis. For example, they would like to manually adjust the color intensities of the matrix and snippets in order to emphasize contrast of sub-regions. The domain experts also expressed desire to pick and search patterns manually in the matrix view, for example, as a means to supplement results of a pattern detection algorithm. Yet, this will require image-based pattern detection algorithms similar to Magnostics [7] but on a much larger scale and in an interactive fashion. Some domain experts mentioned that integrating visualizations of other genomic features into snippets would further assist in finding correlations.

In this work, we focused on exploring squared snippets with a specific pattern (e.g., *dots*). We want to extend HiPiler to support arbitrarily sized snippets of equal ratio, e.g., domains (Sect. 2.3), which requires first investigating appropriate methods for aggregating snippets of different sizes without destroying patterns in the data. Also, we want to provide ways to efficiently show a user’s exploration history and to support collaborative scenarios [21] as mentioned by P1, P3, and P4 during the interviews. Eventually, we want to visually summarize and aggregate data attributes of piles that will enable experts to more seamlessly transition from context-driven (interaction matrix) to knowledge-driven (data attributes) pattern exploration.

### Combining the Matrix and Snippets Approach

Navigable matrices and snippet exploration are complementary approaches. HiPiler integrates both and loosely couples them through brushing and linking. The matrix presents the complete and high-level overview of the data and can show the context for individual ROIs/snippets. However, as discussed in the introduction, that large matrix is a poor means to search for and to visually explore and compare ROIs. Snippet exploration on the other side provides a focused view on the ROIs as well as their visual and data-related features. Future work should try to integrate the best of both worlds; improving the ability to relate between snippets and their context within the matrix while allowing for a snippet-focused exploration. Table 1 summarizes the respective conceptual advantages and disadvantages of both complimentary approaches.

**Table 1.** Strengths (pros) and weaknesses (cons) of the *matrix* and *snippet* approach.

|  | Matrix | Snippets |
| --- | --- | --- |
| **Pros** | Shows ROI in context<br>The neighborhood of patterns can spark new exploration. | High numbers of (far-off) ROIs are comparable<br>Freely arrange and group ROIs according to respective features.<br>Ability to visualize data about sets of ROIs. |
| **Cons** | Impossible to compare spatially far-off ROIs<br>Multiresolution requires manual pan and zoom | Matrix ordering needs to be fixed during exploration.<br>ROIs not visualized in context. |

### Generalizability

Our snippets approach for interactive exploration of many ROIs in a matrix is not limited to genome interaction matrices. It can be applied to any large dataset that can be represented as a matrix and that exhibits a large number of regions of interest and recurring patterns of interest. Most similar to genome interaction matrices are other similarity matrices, e.g., for showing temporal evolution in datasets with specific patterns for temporal change [5]. Another application example are networks with thousands of nodes, represented as adjacency matrices, and which are found in biology (e.g., gene regulatory or protein interaction networks), social application (e.g., Facebook), or computer science (e.g., server networks). ROIs in adjacency matrices can represent topological cliques and clusters, subgraphs, or specific graph motifs resulting in specific visual patterns in the matrix [7]. Adapting snippet exploration to networks requires an appropriate matrix ordering [6] to create visual patterns as well as pattern extraction methods specific to network. These can be topological cluster and motif detection algorithms or visual pattern recognition methods. Another potential application domain for snippet-based exploration are gigapixel images; high content microscopy screening produces very large images of cell cultures or tissues and astronomers study high resolution pictures of galaxies. In these cases analysts are also searching for recurrent visual patterns (cells, galaxies) that are relatively small compared to the entire image.

We are confident that the tasks identified for this project and described in Sect. 2.4, do generalize well to these areas (Sect. 4.1). The bold-faced titles of T1-T6 describe generic tasks while the reasoning for each task depends on the specifics of the data type that is to be explored.

### Scalability and Limitations

The current version of HiPiler can handle up to 2000 snippets, while simultaneously showing the large navigable genome interaction matrix. The main limitation is currently the browser cache used for storing snippets and the user interaction history, while graphics are implemented using WebGL shaders. Future versions of HiPiler can improve scalability by moving parts of the application logic and snippet caching to the back-end. It is important to point out that the domain experts had mentioned in the interviews that they will likely never explore more than a thousand snippets at a time. Yet, the concept behind HiPiler—interactively exploring snippets—is scalable to larger numbers of snippets as snippets can be filtered, and aggregated into piles.

Finally, it is not yet clear how beneficial the piling metaphor would be for snippets of different aspect ratios in terms of aggregation through piling. While it is technically straightforward to visually scale patterns to equal size and aspect ratios, domain experts are not sure what the aggregation of differently sized patterns would mean biologically as this would lead to many non-trivial normalization issues. Also, some biological features, e.g., checkerboard pattern (Sect. 2.3), result in patterns too large to be visualized and explored as snippets. Visualization of such large-scale patterns requires further specialized visualization techniques.

## 9 CONCLUSION

We have introduced HiPiler—a visual interface that enables the exploration of large genome interaction matrices based on many small ROIs through interactive small multiples. In a user study we found that our proposed snippets approach meets the needs of domain experts and complements existing heatmap-based approaches. The tasks identified in our interviews (see Sect. 2.4) prove to be a valid basis for the design of the HiPiler interface and visualizations. Based on our experience with analysis tasks for other large matrices and image data, as well as the successful evaluation of our approach in the user study, we conclude that HiPiler is very likely generalizable and could be applied to other application domains.

We found that snippet-based exploration is an efficient means to explore local patterns in very large matrices. Genome scientists described our tool as highly useful, appropriate, as well as understandable and they anticipate a positive impact on their research. Removing the context of the ROIs did not hinder the explorability if the principal focus is on localized patterns. We showed that the ability to aggregate several snippets into piles reduces visual complexity and aids highlighting pattern diversity. The snippets approach can provide new context to the ROIs, which is useful for pattern exploration along external measures or metadata, beyond the domain of genome interaction matrices.

## ACKNOWLEDGMENTS

The authors wish to thank N. Abdennur, B. Alver, H. Belaghzal, A. van den Berg, J. Dekker, G. Fudenberg, J. Gibcus, A. Goloborodko, D. Gorkin, M. Imakaev, Y. Liu, L. Mirny, J. Nübler, P. Park, H. Strobelt, and S. Wang. This work was supported in part by the National Institutes of Health (U01 CA200059 and R00 HG007583).

